# Humans prioritize walking efficiency or walking stability based on environmental risk

**DOI:** 10.1101/2022.03.13.482180

**Authors:** Ashwini Kulkarni, Chuyi Cui, Shirley Rietdyk, Satyajit Ambike

## Abstract

In human gait, the body’s mechanical energy at the end of one step is reused to achieve forward progression during the subsequent step, thereby reducing the required muscle work. During the single stance phase, humans rely on the largely uncontrolled passive inverted pendular motion of the body to perpetuate forward motion. These passive body dynamics, while improving walking efficiency, also indicate that lower passive dynamic stability in the anterior direction since the individual will be less able to withstand a forward external perturbation. Here we test the novel hypothesis that humans manipulate passive anterior-posterior (AP) stability via active selection of step length to either achieve energy-efficient gait or to improve stability when it is threatened. We computed the AP margin of stability, which quantifies the passive dynamic stability of gait, for multiple steps as healthy young adults (N=20) walked on a clear and on an obstructed walkway. Participants used passive dynamics to achieve energy-efficient gait for all but one step; when crossing the obstacle with the leading limb, AP margin of stability was increased. This increase indicated caution to offset the greater risk of falling after a potential trip. Furthermore, AP margin of stability increased while approaching the obstacle, indicating that humans proactively manipulate the passive dynamics to meet the demands of the locomotor task. Finally, the step length and the center of mass motion co-varied to maintain the AP margin of stability for all steps in both tasks at the specific values for each step. We conclude that humans actively regulate step length to maintain specific levels of passive dynamic stability for each step during unobstructed and obstructed gait.

## 1. Introduction

Gymnasts exploit the mechanical properties of their bodies to achieve remarkable feats. While performing a triple axel, for example, an ice-skater holds her arms close to or farther from her body to modulate her angular velocity. High velocity during the flight phase allows for more turns [1], whereas lower velocity close to landing improves safety [2].

The ability and propensity to adapt the body’s mechanical properties to facilitate motion is not limited to trained sportspersons; it is a feature of the walking patterns of most adults. On clear and level walkways, adults use a characteristic step length to recycle the kinetic energy at the end of a step to rotate the body over the new stance foot. Part of the forward progression of the body is thus achieved passively, which improves walking efficiency [3, 4]. However, if the environment poses a threat to stability, the passive forward motion can be altered to improve gait stability rather than efficiency. Such adaptation is suggested by the difference in the behaviors of young and older adults. While stepping over an obstacle, older adults rely less on the passive forward motion compared to young adults [5], since age-related declines in strength and coordination make it more difficult to recover from a trip. In other words, since, after a trip, forward passive motion makes a forward fall more likely, older adults proactively improve their passive stability by reducing this motion while crossing obstacles.

This tradeoff between stability and efficiency has been supported by the study of the margin of stability in the anterior-posterior direction (MOS_AP_; abbreviations in Table 1). MOS_AP_ has been used to identify differences in the recruitment of passive dynamics across age groups [5–9], in pathological populations [10–12], and in dual tasks [13]. MOS_AP_, based on the inverted pendulum model of gait, and usually computed at heel contact, is the distance from the anterior boundary of the base of support (BOS; the leading heel) to the extrapolated center of mass (XcoM), which reflects the center of mass (CoM) state [14, 15]. The condition XcoM ahead of the BOS boundary at heel contact indicates efficient gait: the body has sufficient kinetic energy to passively rock over the new stance ankle. More anterior location of XcoM indicates greater forward passive motion and greater efficiency. At the same time, a forward perturbation during the swing phase would accentuate the greater forward passive motion, making a forward fall more likely. Therefore, a more anterior location of the XcoM at heel contact is also interpreted as lower passive dynamic stability in the anterior direction [5, 13].

**Table 1.** Abbreviations.

|  |  |
| --- | --- |
| Anterior-posterior | AP |
| Medial-lateral | ML |
| Foot placements | fp |
| Center of mass | CoM |
| Center of mass velocity | V<sub>CoM</sub> |
| Leg length | $l$ |
| Acceleration due to gravity | $g$ |
| Base of support | BOS |
| Extrapolated center of mass | XcoM |
| Margin of stability in anterior-posterior direction | MOS<sub>AP</sub> |
| Uncontrolled manifold | UCM |
| Orthogonal manifold | ORT |
| Synergy index | $\Delta V$ |
| Synergy index z transformed | $\Delta V_z$ |
| Variance along the uncontrolled manifold | $V_{UCM}$ |
| Variance along the orthogonal manifold | $V_{ORT}$ |

Here, we advance in three ways our understanding of the role of passive dynamics during gait based on MOS_AP_. First, we demonstrate that adaptation in passive dynamics in response to environmental threats to stability is apparent in young adults as well. We show that young adults increase passive dynamic stability while crossing an obstacle compared to unobstructed walking. Second, we demonstrate that the shift in passive dynamic stability occurs in the steps leading up to the obstacle. This novel finding is consistent with the notion that motor behavior usually does not change instantaneously, but evolves over characteristic times determined in part by the inertia of the body [16]. Third, we demonstrate that humans modulate step length to control MOS_AP_. We show that MOS_AP_ is similar for the steps of unobstructed gait, and it is maintained at different values for specific steps while approaching and crossing an obstacle. Hof [15] showed that a consistent MOS_AP_ yields a stable walking cycle in a mathematical model, and hypothesized that humans would benefit from this strategy as well [17]. It has been suggested that the variability rather than the average value of MOS better represents passive dynamic stability of human gait [18]. In the anterior-posterior (AP) direction specifically, indirect evidence for this hypothesis is provided by the fact that persons with Multiple Sclerosis (MS) exhibit greater MOS_AP_ variability compared to healthy controls [11], and within the group of persons with MS, fallers show greater MOS_AP_ variability than non-fallers [10].

We sought direct evidence for the control of MOS_AP_ in the unobstructed and obstructed gait of healthy young adults. Since the MOS_AP_ is a function of step length and XcoM, we hypothesized that the central nervous system responds to changes the XcoM with a corresponding correction in step length so that the MOS_AP_ itself is relatively invariant at each heel contact. However, the specific values of MOS_AP_ that are maintained via the covariation in XcoM and step length are different at various steps during obstructed gait.

We employed the uncontrolled manifold (UCM) method [19] to test this hypothesis. For each heel contact, we computed the synergy index that quantifies the covariation between XcoM and step length. A positive synergy index indicates that MOS_AP_ was actively controlled, i.e., stabilized at the specific across-trial mean MOS_AP_ for that step. A higher value indicates a stronger synergy or higher stability of MOS_AP_. We emphasize that stability of MOS_AP_ is different from the stability of gait. MOS_AP_ is the measure of *passive* dynamic gait stability. In contrast, the synergy index indicates the efficacy of *active* control at stabilizing or maintaining MOS_AP_ at a specific value for a given step.

We computed MOS_AP_ and the synergy index for multiple steps for unobstructed gait trials and for trials where participants approached an obstacle, stepped over it, and continued walking till the end of the walkway. We hypothesized a task by step interaction for MOS_AP_ (H1). The MOS_AP_ will not be different across tasks (obstructed and unobstructed) for the steps at the start and end of the walkway, but the MOS_AP_ will be higher for the approach steps 1-2 steps before the obstacle and for the crossing steps. These changes will indicate prioritization of safety over efficiency [5, 6, 20]. Next, we hypothesized that the synergy index will be greater than zero, indicating that the step length and the XcoM co-vary to stabilize MOS_AP_ for all steps in both tasks (H2). Finally, we hypothesized a task by step interaction for the synergy index (H3). The synergy index will not be different across tasks for the steps at the start and end of the walkway, but the synergy index will be lower while crossing an obstacle placed in the middle of the walkway. The lower synergy index will reflect the greater motor demands associated with the crossing steps; larger muscle activations required for stepping over the obstacle [21] will be associated with greater noise [22], which will make stabilization more difficult.

## 2. Materials and Methods

### 2.1 Participants

Twenty-six healthy young adults participated in the study. We excluded six participants due to poor kinematic tracking. Data from 20 participants (14 women, 22.3 ± 3.7 years, 1.7 ± 0.1 m, 66.9 ± 14.6 kg) were used for analysis. All participants walked without aid, had no orthopedic, neuromuscular, or dementia disorders, and were independent in daily activities. Vision was normal or corrected-to-normal. The study was approved by Purdue University’s Institutional Review Board, and all participants provided written informed consent (Protocol number: IRB-2021-331).

### 2.2 Equipment and Procedures

We assessed leg dominance using the Waterloo Footedness Questionnaire— Revised [23]. Participants walked at their self-selected speed on a 6.0 m walkway and stepped over an obstacle when present (Fig. 1A). The obstacle was 100 cm wide × 0.4 cm deep. The height of the obstacle was scaled to 25% of the participant’s leg length. The obstacle was made of black Masonite and designed to tip if contacted. The starting position was determined for each participant such that they took five steps before reaching the obstacle, crossed the obstacle naturally with the right leg first and stopped three to four steps later (Fig. 1A).

**Figure 1.**
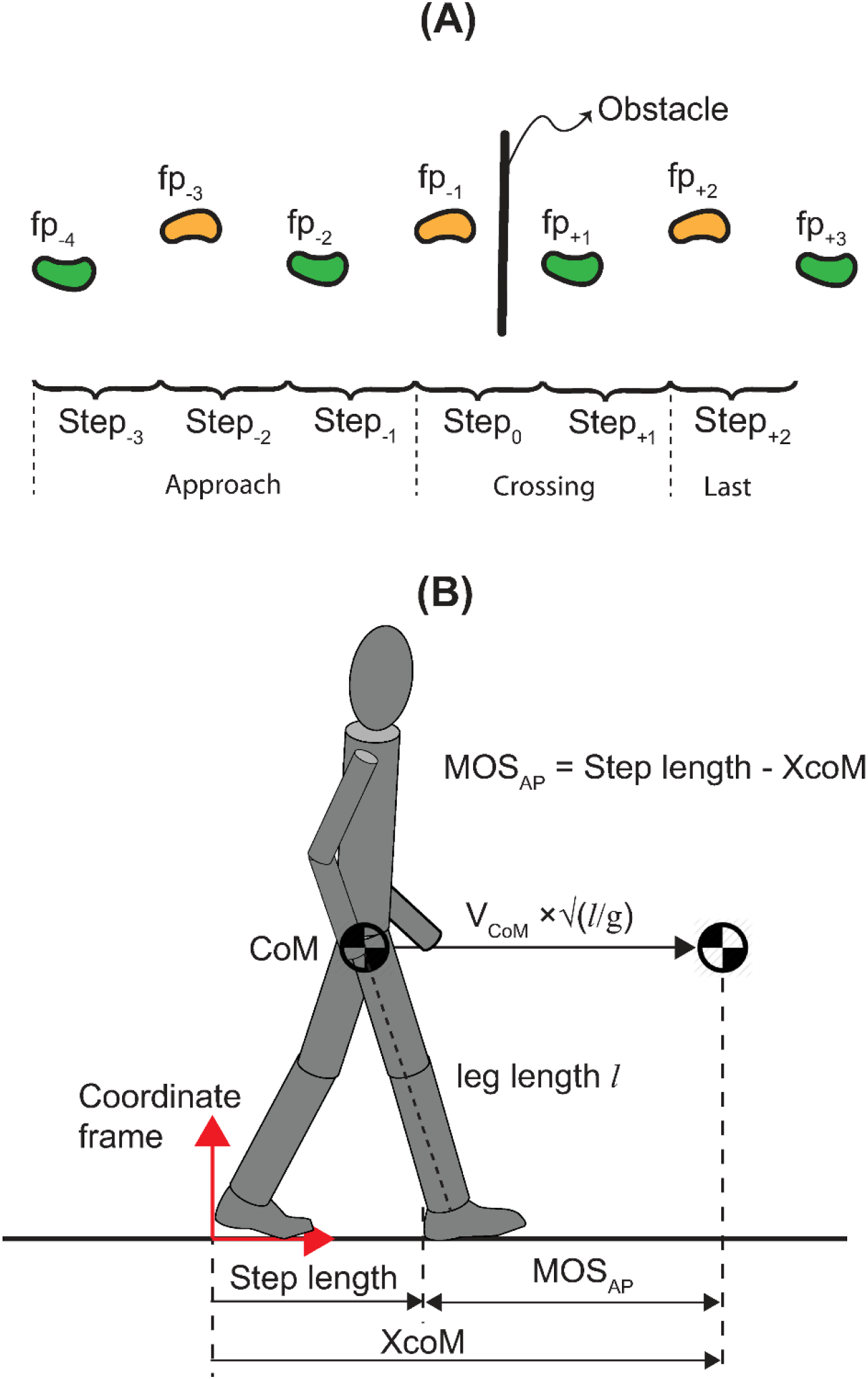
Experimental task and basic definitions. Illustration of foot placements while approaching (fp_-4_ to fp_-1_), crossing (fp_+1_ and fp_+2_) and after crossing (fp_+3_) the obstacle, and of steps while approaching (Step_-3_ to Step_-1_), crossing (Step_0_ and Step_+1_) and after crossing (Step_+2_) the obstacle **(A)**. Definitions of the extrapolated center of mass (XcoM) and margin of stability, MOS_AP_. Step length, XcoM and MOS_AP_ are computed at the moment of lead heel contact in a coordinate frame located where the rear heel contacted the ground **(B)**.

Participants first performed 20 trials without an obstacle (no obstacle task), followed by 20 trials of walking with an obstacle (obstacle-crossing task). We collected kinematic data at 100 Hz with a motion capture system (Vicon Vero, Oxford, UK) with marker clusters placed bilaterally on the lower back, thigh, shank, and foot. We digitized the joint centers and posterior aspect of the heels to identify their locations relative to the marker clusters. We also digitized the top edge of the obstacle to identify its position.

### 2.3 Analysis

Across all trials and all participants, the obstacle was contacted and tipped 12 times out of 3360 trials (0.4 % of trials). We did not include the contact trials in our analysis. Furthermore, some trials were discarded due to poor kinematic tracking. To have the same number of trials for all participants and tasks, we selected 15 trials with good kinematic data. Fifteen trials are sufficient for reliable quantification of the synergy variables [24]. We filtered all kinematic data using a zero-lag, 4^th^ order, low-pass Butterworth filter with a cut-off of 7 Hz. We identified seven foot placements (Fig. 1A) using the AP position of the heel [25]. We quantified spatiotemporal gait parameters and margin of stability at heel contact at the seven foot placements (fp_-4_ to fp_+3_) and six steps (Step_-3_ to Step_+2_; Fig. 1A). Step length was defined as the distance between two consecutive heel contacts. CoM position was computed as the centroid of the triangle formed by the left and right anterior superior iliac spines and the center of the left and right posterior superior iliac spines [26]. CoM velocity was obtained by differentiating the CoM position data. The extrapolated center of mass (XcoM) was calculated in the sagittal plane as [14]:

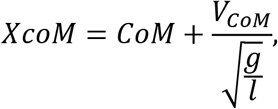

where CoM is the anterior-posterior CoM position, V_CoM_ is the anterior-posterior CoM velocity, *g* is the acceleration due to gravity, and *l* is the participant’s leg length (Fig. 1B). Leg length was calculated as the sagittal-plane distance between the CoM and the ankle of the limb that contacted the ground. We used the average of the leg length values obtained from the 15 trials for each step to compute the XcoM for that step [18]. We computed MOS_AP_ at the instant of leading heel contact in a coordinate frame fixed at the location of the rear heel contact (Fig. 1B):

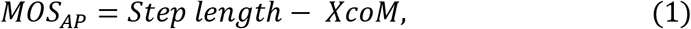

i.e., the distance from the anterior boundary of the BOS, defined by the position of the leading heel, to the XcoM. Negative MOS_AP_ (XcoM ahead of the anterior BOS boundary) at heel contact indicates that the body possesses sufficient energy to passively rotate beyond the upright position and fall forward (assuming that the energy loss at heel contact and the energy input during the subsequent push-off are either equal or negligible). Conversely, positive MOS_AP_ (XcoM is behind the anterior BOS boundary) indicates that the body cannot passively rotate beyond the upright position, and it will eventually fall backward.

For a passive walker, MOS_AP_ = 0 serves as a threshold for detecting unstable gait. Hof demonstrated that stable gait arises with XcoM ahead of the BOS (MOS_AP_ < 0; Eqn. 1) in a mathematical model of a passive walker [15]. Conversely, MOS_AP_ > 0 would mean that the forward progression of the passive walker will stop, and the gait will be unstable.

In this work, we are interested in the forward loss of stability arising from a trip. Therefore, we invert the interpretation of the MOS_AP_ from that for the passive walker. We consider MOS_AP_ directly proportional to gait stability in the anterior direction. That is, lower MOS_AP_ (more anterior location of the XcoM) indicates less stable gait, since a forward fall is more likely if a perturbation like a trip occurs. Conversely, higher MOS_AP_ (more posterior location of XcoM) indicates more stable gait [13, 27]. Furthermore, MOS_AP_ is inversely proportional to efficiency: lower MOS_AP_ indicates greater passive forward motion, and hence less active propulsion is required to maintain forward progression, and vice-versa. In this way, MOS_AP_ reflects the tradeoff between stability and efficiency.

We used the uncontrolled manifold (UCM) analysis to quantify the synergy stabilizing MOS_AP_ at heel contact. A *synergy* is co-variation in a redundant sets of input body variables that maintains important output variables that define task performance. The UCM method has been widely used to quantify the synergistic covariation in body variables in a variety of human movements including gait ([28, 29]; see [30, 31] for recent reviews). Importantly, in addition to identifying task-specific covariation, the UCM method identifies the salient task variables controlled by the nervous system.

Here, we use the UCM method to evaluate the hypothesis that the input variables – step length and XcoM, co-vary to stabilize the output variable, i.e., MOS_AP_. We performed the analysis separately for each step for both tasks. We first obtain from the constraint equation (Eqn. 1), the Jacobian matrix that relates small changes in the step length and XcoM to changes in MOS_AP_: J = [1 −1]. The one-dimensional null space of this Jacobian defines the UCM, and its one-dimensional compliment defines the orthogonal (ORT) manifold. We pool the across-trial step length and XcoM data for a particular step. The deviation in the step length and XcoM data for each trial from the across-trial mean is projected onto the UCM and the ORT manifolds. The variances in these projections are the V_UCM_ and the V_ORT_, respectively. These variance components yield the synergy index:

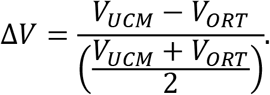

The synergy index ΔV has a threshold value of zero. When ΔV > 0, V_UCM_ > V_ORT_. In general, this implies that the control is organized so that most of the variability in the inputs is channeled along the UCM, and therefore, it does not alter task performance, i.e., the output. Here, ΔV > 0 implies that the step length and XcoM covary to stabilize MOS_AP_. Conversely, when ΔV < 0, V_UCM_ < V_ORT_. This implies that control is organized so that most of the variability in the inputs alters task performance (i.e., change MOS_AP_). In both these cases, when the synergy index differs from zero, we would conclude that MOS_AP_ is a controlled variable. Finally, ΔV = 0 indicates that there is no task-specific co-variation in the step length and XcoM. This result would indicate that MOS_AP_ is not a controlled variable.

The synergy index ΔV ranges from −2 to 2. Therefore, it was z-transformed for statistical analysis [29, 32]:

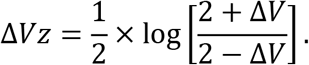

Note that ΔV = 0 translates to ΔVz = 0. We report ΔVz values in the Results section and use ΔVz values to draw inferences, consistent with most previous studies [30].

### 2.4 Statistical analysis

To determine whether the gait task and foot placement affected MOS_AP_ (H1), we performed a two-way (task × step) repeated measures ANOVA. To identify the source of changes in MOS_AP_, we performed the two-way ANOVA separately on the CoM position, CoM velocity at heel contact and step length. To determine whether the synergies were present, we performed separate, one-sample t-tests to test if ΔVz was significantly different from zero for each step in the two tasks (H2). To determine whether the gait task and foot placement affected the synergy index (H3), we performed a task × step repeated-measures ANOVA. To identify the source of changes in the synergy index, we performed the two-way ANOVA separately on the variance components (V_UCM_, V_ORT_). For all ANOVA tests, we fit a generalized linear model with random effects. Recall that the UCM analysis utilizes data from all 15 trials to yield a single value each for the synergy index, V_UCM_, and V_ORT_. Therefore, participant was the random effect for these ANOVAs. However, data from all 15 trials was used to analyze all the remaining variables. Therefore, the trial number within each participant was the nested random effect for these ANOVAs. Tukey-Kramer adjustments were used to perform the following planned pairwise comparisons: (1) across-task comparison at each of the six steps (or seven foot placements), and (2) all across-step (or foot placement) comparisons for the obstacle-crossing task only. All analyses were performed using the PROC GLIMMIX procedure in SAS 9.4 (Cary, NC, USA) with significance set at 0.05.

## 3. Results

The Waterloo Footedness Questionnaire scores were 8 + 5, indicating that all participants were right-leg dominant.

Next, we use data from one participant (Fig. 2) to outline the overall results. The detailed statistical results are presented in the following subsections. Figure 2 illustrates (1) the changes in various outcome variables while approaching, crossing, and resuming gait after crossing an obstacle, and (2) how the variables compare with unobstructed gait for each step. All outcome variables showed a task × step interaction (supporting H1 and H3) indicating that the pattern of changes in the variables across steps was different for the obstacle-crossing task compared to the no obstacle task. The largest across-task changes were observed in all outcomes for the two crossing steps (Step_0_ and Step_+1_), and across-task changes were apparent in some outcomes in the approach to the obstacle (Step_-3_ to Step_-1_). All variables showed across-step changes for the obstacle-crossing task. The synergy index was greater than zero, indicating that the step length and the XcoM co-varied to stabilize MOS_AP_ for all steps in both tasks (supporting H2). In the detailed results below, we first present the results for MOS_AP_, followed by results for the variables that constitute MOS_AP_: CoM position relative to rear heel, CoM velocity at heel contact, and step length. We then present the results for the UCM outcome variables quantifying the stability of MOS_AP_.

**Figure 2.**
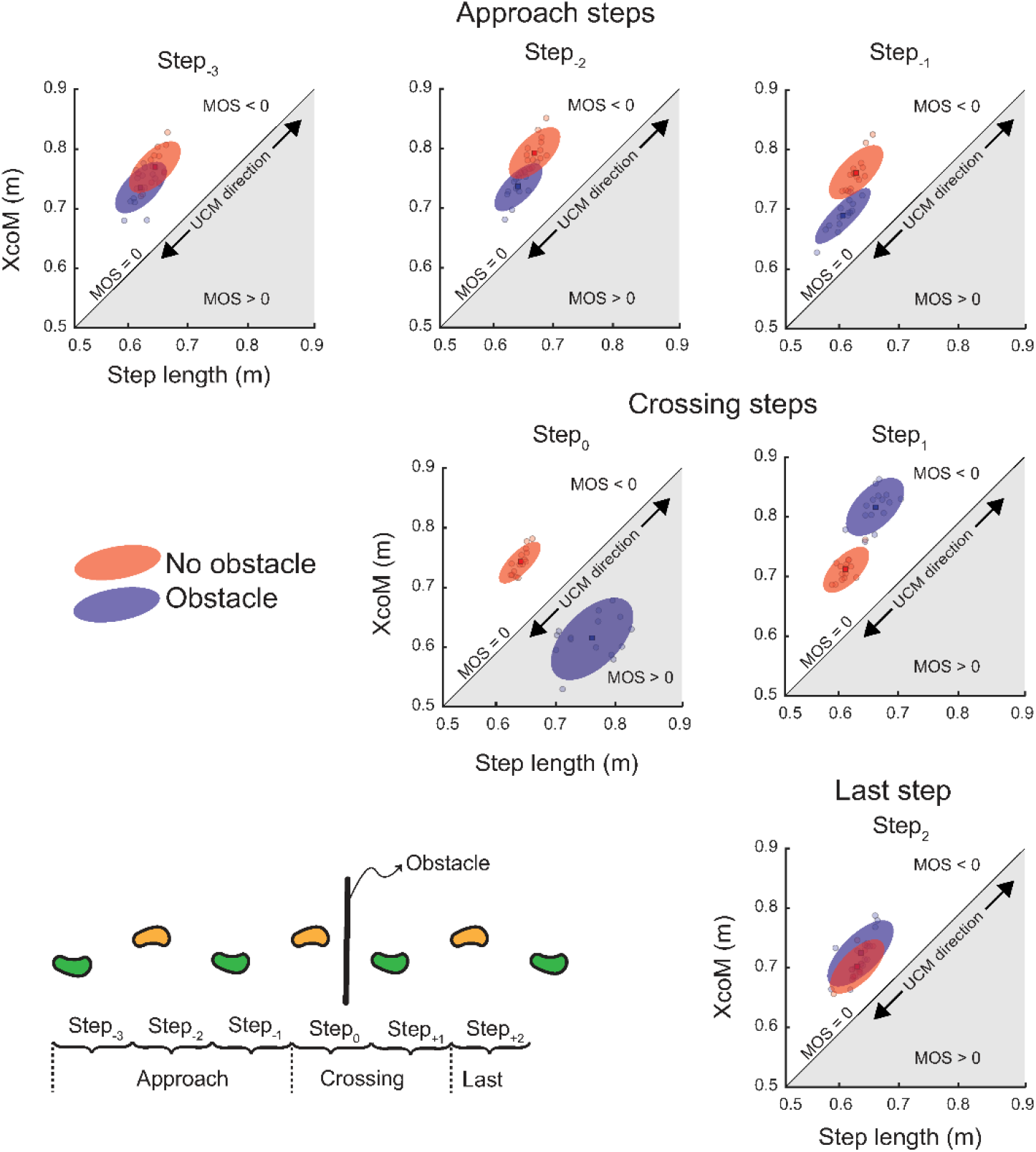
Representative data from one participant. Figure illustrates changes in various outcomes while approaching and crossing an obstacle, and then resuming unobstructed gait (blue ellipses). Steps during unobstructed gait are included for comparison (red ellipses). The 45° line separates the regions of positive and negative MOS_AP_; the white region above the line represents passively unstable behavior, and the gray region below the line represents passively stable behavior. The 45° line also represents the direction of the uncontrolled manifold (UCM).

Variations in step length and XcoM along this direction do not change MOS_AP_. The ellipses are centered at the centroid of the across-trial data for each step, and the major axes are aligned with the 45° line. However, the flatness of the ellipses is representative. Changes in mean values of the MOS_AP_ are reflected in the position of the ellipses in each chart. The flatness of the ellipse reflects the synergy index, with flatter ellipses indicating higher synergy indices that reflect stronger covariation between the step length and XcoM.

### 3.1 Margin of stability

#### MOS_AP_

We observed a task × step interaction for MOS_AP_ (F_6,4167_ = 886.89, p < 0.001; η_p_^2^ = 0.9; Fig. 3A). Post-hoc comparisons across tasks revealed that MOS_AP_ was higher (more stable) for the obstacle-crossing task than the no obstacle task at all except the first (fp_-4_) and last two foot placements (fp_+2_ and fp_+3_). MOS_AP_ was lower (less stable) for fp_+2_ (after trail foot crossing) for the obstacle-crossing task. All pair-wise across-step comparisons for the obstacle-crossing task are depicted in Fig. 3A. For brevity, we describe only some of the significant differences. Changes in MOS_AP_ occurred before the obstacle was reached; MOS_AP_ was higher (more stable) for fp_-1_ relative to the two preceding foot placements. For the two crossing steps, MOS_AP_ first increased to the highest value for fp_+1_, and then decreased to the lowest value for fp_+2_ across all foot placements.

**Figure 3.**
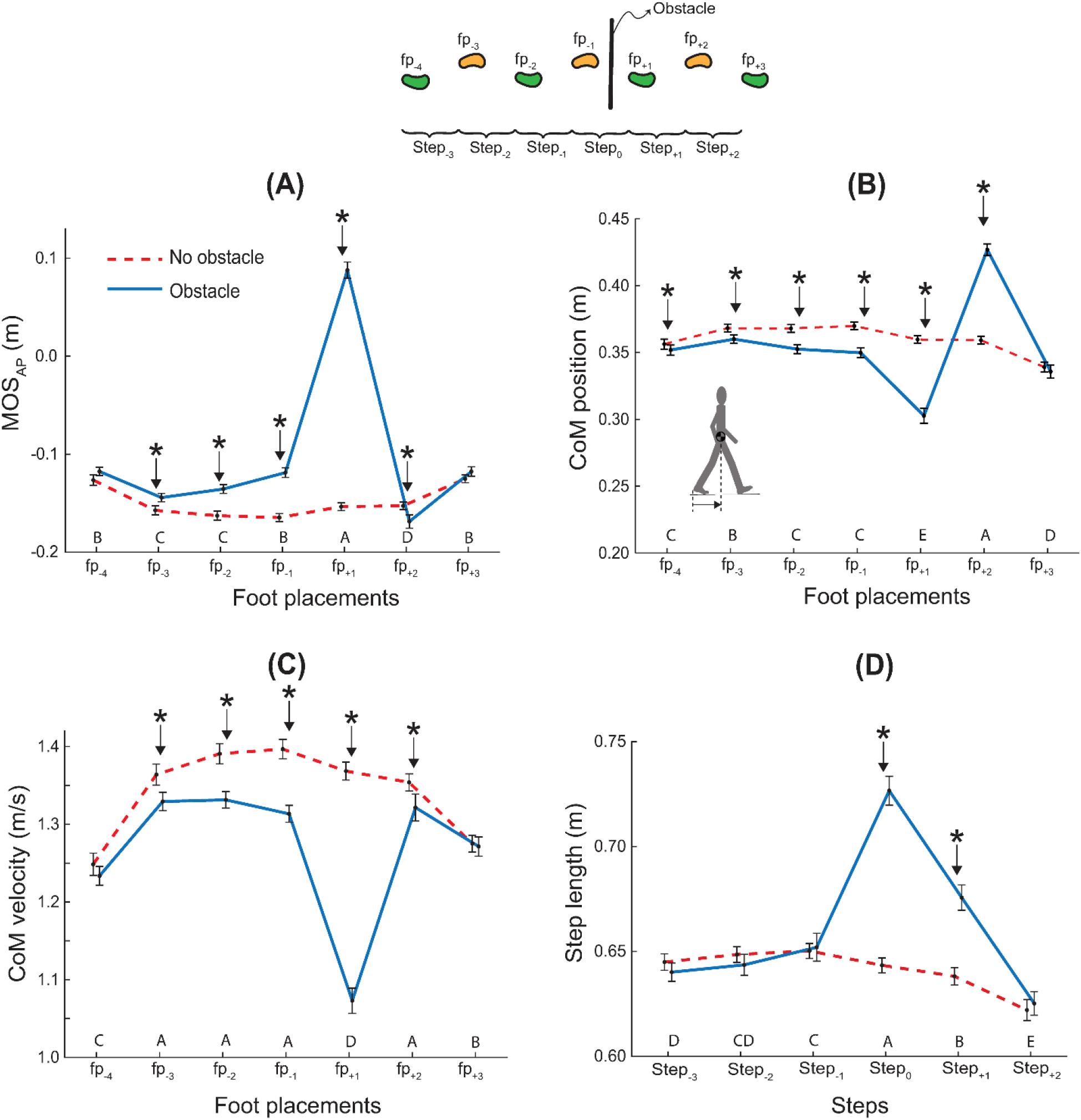
MOS_AP_ and its components. MOS_AP_ **(A)**, CoM position at heel contact **(B)**, CoM velocity at heel contact **(C)** and step length **(D)**. Data are across-subject means (N=20), and error bars denote standard error. Asterisks denote significant task difference at that step (p < 0.05). Across-step pairwise comparisons for the obstacle-crossing task are indicated by the row of letters above the horizontal axis for each panel. Steps that do not have letters in common are significantly different from each other (i.e., A is different from B, but A is not different from AB).

#### CoM position relative to the rear heel

A task × step interaction was observed for the CoM position relative to the rear heel (F_6,4167_ = 391.20, p < 0.001; η_p_^2^ = 0.9; Fig. 3B). Post-hoc comparisons across tasks revealed that CoM was closer to the rear heel at the foot placements of all approach steps and the lead crossing step (fp_-4_ to fp_+1_) for the obstacle-crossing task than the no obstacle task. Conversely, CoM was closer to the front heel at the foot placement for the trail crossing step (fp_+2_) for the obstacle-crossing task. All pair-wise across-step comparisons for the obstacle-crossing task are depicted in Fig. 3B. For brevity, we describe only some of the significant differences. CoM position changed for the last three foot placements. The CoM was closer to the rear foot at the foot placement for the lead crossing step (fp_+1_). It shifted forward, so that it was closer to the front heel at the foot placement for the trail crossing step (fp_+2_). Finally, for the last step (fp_+3_), CoM position was consistent with that for the approach steps.

#### CoM velocity at heel contact

We observed a task × step interaction for the CoM velocity at heel contact (F_6,4167_= 161.52, p < 0.001; η_p_^2^ = 0.8; Fig. 3C). Post-hoc comparisons across tasks revealed that the CoM velocity was lower for the obstacle-crossing task compared to the no obstacle task at all but the first and the last foot placements (fp_-4_ and fp_+3_). Post-hoc comparisons across steps for the obstacle-crossing task revealed that the CoM velocity was lower at the first (fp_-4_) and last (fp_+3_) foot placement compared to other foot placements, except at the lead foot crossing (fp_+1_), where the CoM velocity was lower compared to all other foot placements.

#### Step length

We observed a task × step interaction for the step length (F_5,3569_ = 149.09; p < 0.001; η_p_^2^ = 0.9; Fig. 3D). Post-hoc comparisons across tasks revealed that step length was longer during the obstacle-crossing task compared to the no obstacle task at the lead (step_0_) and trail crossing (step_+1_) steps. Post-hoc comparisons across steps for the obstacle-crossing task revealed that step length was higher for step_-1_ compared to step_-3_. That is, relative to the early step, the step length increased one step before the first crossing step. Step length was highest for the lead crossing step (step_0_), and then it shortened for the trail crossing step (step_+1_), but remained longer than all other steps. Finally, step length was the shortest for the last step (Step_+2_) compared to all other steps.

### 3.2 UCM variables

#### Synergy index

The synergy index (ΔVz) was significantly greater than zero at all steps for both tasks (t_19_ < 15.97, p < 0.01; Fig. 4A).

**Figure 4.**
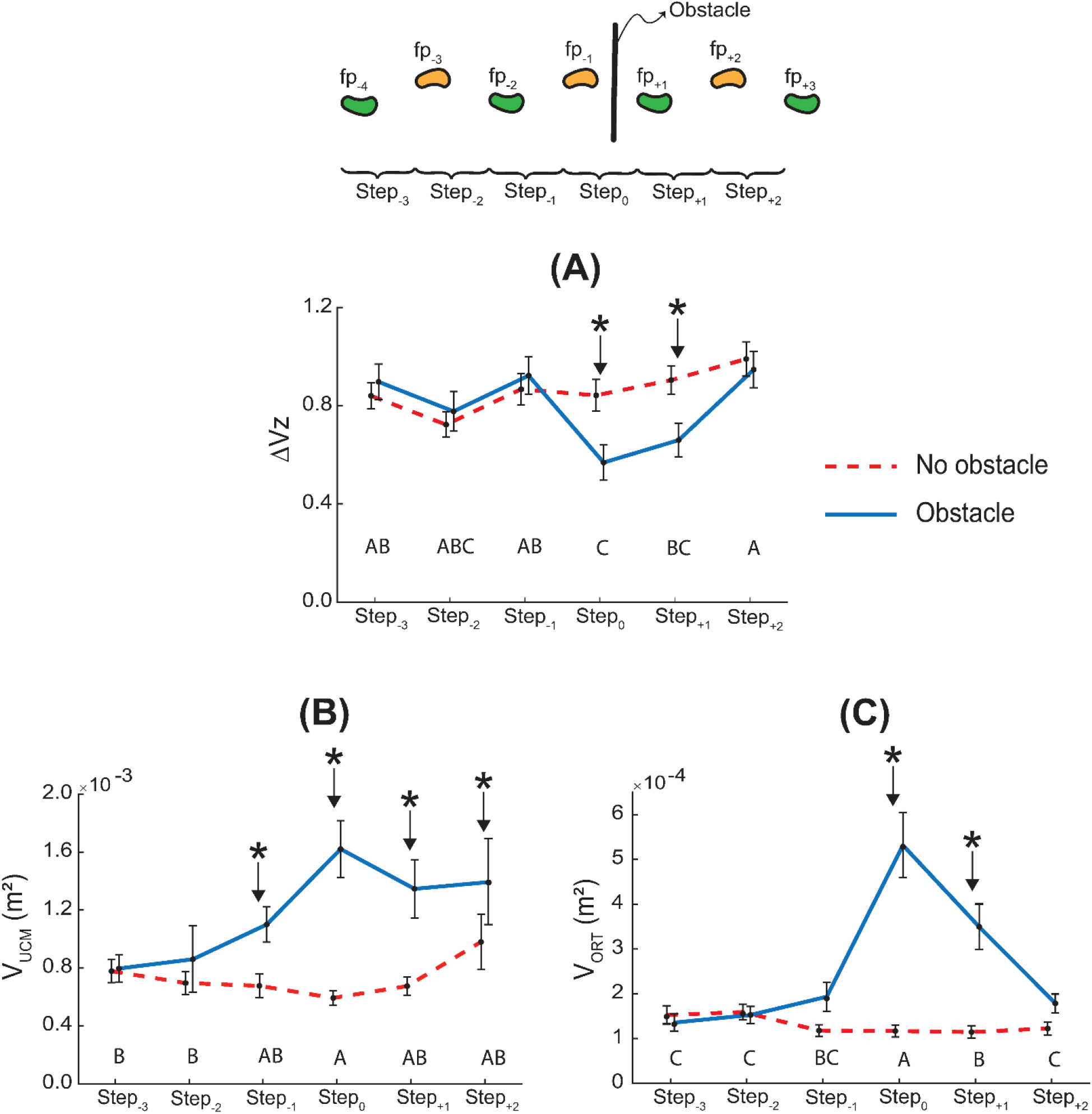
Outcomes of the synergy analysis. Synergy index **(A)**, V_UCM_ **(B)**, and V_ORT_ **(C)**. Data are across-subject means (N=20), and error bars denote standard error. Asterisks denote significant across-task differences at that step (p < 0.05). Across-step pairwise comparisons for the obstacle-crossing task are indicated by the row of letters above the horizontal axis for each panel. Steps that do not have letters in common are significantly different from each other (i.e., A is different from B, but A is not different from AB).

There was a task × step interaction for the synergy index ΔVz (F_5,209_ = 2.89, p = 0.015; η_p_^2^ = 0.3; Fig. 4A). Post-hoc comparisons across tasks revealed that ΔVz was lower during the obstacle-crossing task compared to the no obstacle task for only the lead (step_0_) and trail crossing steps (step+1). Post-hoc comparisons across steps for the obstacle-crossing task revealed that ΔVz was lower at the lead crossing step (step_0_) compared to all non-crossing steps except Step_-2_, and ΔVz was lower at the trail crossing step (step+1) compared to the last step (Step_+2_).

#### Variance components

There was a task × step interaction for V_UCM_ (F_5,209_ = 3.12, p = 0.009; η_p_^2^ = 0.3; Fig. 4B). Post-hoc comparisons across tasks revealed that V_UCM_ was higher for the obstacle-crossing task compared to the no obstacle task at all but the first two steps (Step_-3_ and Step_-2_). Post-hoc comparisons across steps for the obstacle-crossing task revealed that V_UCM_ was higher at the lead crossing step (step_0_) compared to the first two steps (Step_-3_ and Step_-2_).

There was a task × step interaction for V_ORT_ (F_5,209_ = 16.51, p < 0.001; η_p_^2^ = 0.3; Fig. 4C). Post-hoc comparisons across task revealed that V_ORT_ was higher during the obstacle-crossing task compared to the no obstacle task at the lead (step_0_) and trail (step+1) crossing steps. Post-hoc comparisons across steps for the obstacle-crossing task revealed that V_ORT_ was higher at the lead crossing step (step_0_) compared to all other steps, and V_ORT_ was higher at the trail crossing step (step+1) compared to all steps except the lead crossing step (lower for step+1 compared to step_0_) and the preceding step (step-1; no significant difference).

## 4. Discussion

Our goals were to establish that young adults proactively change MOS_AP_ while approaching and then crossing an obstacle, and that MOS_AP_ is actively controlled during unobstructed as well as obstructed walking. We had hypothesized a task by step interaction for MOS_AP_ (H1). We observed proactive changes in MOS_AP_ in anticipation of a threat to stability; MOS_AP_ changed across steps for unobstructed versus obstructed gait (supporting H1). Next, we had hypothesized that the synergy index will be greater than zero (H2). We report the novel finding that the central nervous system responds to changes in the body’s motion with a corresponding correction in step length so that the MOS_AP_ itself is approximately invariant at each heel contact (supporting H2). Finally, we had hypothesized a task by step interaction for the synergy index (H3). We observed that the synergy index was lower while crossing an obstacle compared to earlier and later steps and compared to the corresponding steps during unobstructed gait (supporting H3). We argue below that (1) the proactive changes in the MOS_AP_ for the obstacle-crossing task reflect a stability-efficiency tradeoff; and (2) the positive synergy indices indicate that the passive dynamic stability is not a byproduct of another process, but that the step length is actively controlled to exploit consistently the passive dynamics to achieve either efficient or more stable gait.

### 4.1 Proactive changes in MOS_AP_ and the compromise between stability and energy efficiency

The first major finding of this study is that the MOS_AP_ changes while approaching the obstacle. When compared to unobstructed gait, greater stability was apparent three steps before the obstacle (Fig. 3A). Changes in MOS_AP_ during approach and crossing resulted from reduced speed, a more posterior CoM, and, for some steps, increased step length (Figs. 3B, 3C, 3D). Previous work has demonstrated that changes in MOS_AP_ and MOS_ML_ were evident during the swing phase immediately prior to taking unnaturally long or quick steps [33], and in MOS_ML_ one step before reaching an obstacle [34]. Here we extend this finding and demonstrate that MOS_AP_ changes several steps before reaching the obstacle, indicating that the transition from unobstructed to obstructed gait occurs over several steps.

Our findings extend the argument that passive dynamic stability is modulated in response to perceived risk [6, 27]. For example, the substantive increase in MOS_AP_ for the lead crossing step (Figs. 2, 3A) reflects cautious gait and may be a preemptive strategy against a potential trip that could lead to a forward fall [12]. Similarly, the even-more-substantive transition back to the least stable passive dynamics (lowest MOS_AP_; Fig. 3A) following the trail crossing step may reflect the reduced risk of a forward fall after the trail foot has crossed the obstacle, and the exploitation of passive dynamics to propel the body forward to regain gait speed.

These fluctuations in passive dynamic stability increase the energetic cost of locomotion. The increase in MOS_AP_ over approach steps, and especially the positive MOS_AP_ for the crossing step, indicates that the trailing leg must push off more so that the body can rotate about and beyond the stance ankle [35]. Furthermore, the lead crossing step (step_0_) is a slower and *longer* step (Figs 3C, 3D), which is inconsistent with the typical combination of a slower and *shorter* step during unobstructed gait which reduces energetic cost [4, 36]. Therefore, our results reflect a tradeoff between stability and energy efficiency, consistent with similar arguments offered in the context of stair descent [6], and model-based optimization computations of obstacle crossing behaviors [20].

In sum, inspecting changes in MOS_AP_ across tasks and steps leads to the conclusion that MOS_AP_ is proactively adjusted (1) during the approach, likely to facilitate the transition from unobstructed gait to the movements required to clear the obstacle, (2) while crossing an obstacle, likely to prioritize safety over energy optimality, and (3) when resuming level gait to regain gait speed. The large fluctuations in MOS_AP_ reflect a tradeoff between stability and energy efficiency.

### 4.2 Active control of MOS_AP_ during unobstructed and obstructed gait The synergy index provides evidence for the control of MOS_AP_

The second major finding of this study is that MOS_AP_ is a controlled variable for obstructed as well as unobstructed gait (ΔVz > 0 for all steps; Fig. 4A). In particular, the synergy index remains significantly larger than zero, even though the input variables that define the MOS_AP_ change over multiple steps for the obstacle-crossing task (Figs 2 and 3). Thus, the UCM analysis provides strong quantitative evidence that MOS_AP_ is controlled.

The variance components (Figs 4B, 4C) provide information regarding the underlying processes that stabilize MOS_AP_. Higher V_ORT_ indicates higher variability in MOS_AP_, whereas higher V_UCM_ indicates greater compensatory covariance between XcoM and step length. We observed that MOS_AP_ is more variable for the crossing steps (up to 205% increase in V_ORT_ compared to earlier steps; Fig. 4C). This likely arises from the larger muscle activations and joint moments required for the crossing steps compared to unobstructed steps [21, 37]. Higher activations would increase signal-dependent noise [22], which will lead to more variable MOS_AP_. However, this increase is offset by an increase in V_UCM_ (up to 103% increase compared to earlier steps; Fig. 4B). This compensation leads to MOS_AP_ stabilization overall (Fig. 4A).

#### Does the synergy arise from passive mechanics, or do neural mechanisms indicate active control?

It is unlikely that passive mechanics of the gait cycle alone can explain the positive synergy indices that we observed. Rather, both passive mechanics and active control are responsible for our data. It is indeed likely that mechanics contribute; for example, a greater push off force would result in a more anterior XcoM at the next heel contact, but it would also tend to produce longer steps [3], thereby helping to maintain MOS_AP_. However, the step length is not entirely determined by a passively swinging leg. Rather, activity in the leg muscles and power at the hip and knee joints indicate active control of the forward swinging limb, which will alter the step length from what a passively swinging limb would yield [38]. Spinal stretch reflex loops have long been implicated in the control of locomotion [39, 40]. Feldman [41] recently elucidated locomotor control based on his theory of referent configurations which incorporates spinal reflexes. The central idea in this theory is that to intentionally change a limb’s position, the nervous system modifies the parameter λ, which is the threshold muscle length at which α-motor neurons are recruited and the stretch reflex is activated. Furthermore, movements are executed by setting the time courses λ(t) for various muscles. Muscle activations arise via the stretch-reflex loop in relation to their deviations from the corresponding λ to drive the muscles (and hence the body) towards its referent configuration. It is plausible that variations in muscle lengths from their reference values, arising from variations in push off forces, will engage spinal stretch reflexes that will alter step length and contribute to the observed MOS_AP_ synergy [42]. This explanation is also consistent with the view that for steady state, level gait, humans rely on spinal feedback for control in the AP direction [43, 44].

The contribution of neural mechanisms to the MOS_AP_ synergy may be even greater for obstructed gait, and in addition to the spinal processes implicated during unobstructed gait, supraspinal processes may contribute to the synergy. We observed large fluctuations in the MOS_AP_ over the approach and crossing steps (174% increase for fp_+1_ over fp_-1_, and a 292% decline for fp_+2_ over fp_+1_; Fig. 3A). Nevertheless, the synergy index, although lower (38% decline for Step_0_ relative Step-1; Fig. 4A), remained positive. The positive synergy index despite large fluctuations in related variables over consecutive steps suggests the involvement of supraspinal mechanisms. It is generally thought that supraspinal mechanisms influence synergies. The most convincing evidence is that synergies in individuals with neurological problems (persons with Parkinson’s disease, stroke survivors, etc.) are altered compared to healthy, age-matched controls (see [31] for review). Specifically, during obstacle crossing, supra-spinal mechanisms may influence the synergy index by modulating the gain on spinal reflexes. Indeed, supraspinal centers are involved in the control of the obstacle crossing steps [21]. Spinal reflex responses in the stance leg flexors are enhanced, likely due to increased activity in the prefrontal cortex, in preparation for swinging the leg over the obstacle. Furthermore, the prefrontal cortical activity remains enhanced (compared to unobstructed walking) while swinging the leg over the obstacle [21]. It is also known that visual information about the obstacle is gathered during approach and used to alter gait characteristics [45, 46]. Gathering and using visual information also implicates higher brain centers in the control of obstructed gait, supporting our view that the MOS_AP_ synergy arises from spinal and supraspinal neural circuits. We note both spinal and supraspinal structures are frequently mentioned as candidate neurophysiological bases of synergies [47], although the specific mechanisms are unknown.

In summary, the synergy values that we observed could arise partially from passive body mechanics. However, active neurophysiological processes at the spinal and supra-spinal level likely contribute to this signal as well, indicating active control of MOS_AP_.

Our ideas parallel previous work indicating that the MOS in the medio-lateral (ML) direction is controlled during level walking [18, 48–53]. The focus of previous research on ML stability – versus AP stability – is consistent with the view that level gait requires more control along the ML direction, and minimal control along the AP direction [43]. The UCM analyses performed here show that the MOS is also controlled in the AP direction by regulating step length – not only in gait tasks that require proactive adaptations, but also during unobstructed gait.

### 4.3 Limitations

As an indicator of the stability of human gait, MOS_AP_ is challenging to interpret [54]. The strict interpretation is that XcoM ahead of the BOS boundary indicates that gait cannot be stopped within a step. This would indicate instability if the task were to stop walking within a step. However, when the task is to continue walking, Hof’s result is more pertinent: the same condition yields a stable gait in a passive walker by ensuring that the walker does not stall [15].

The challenge in interpreting MOS_AP_ for human gait arise when extending this logic to (1) different adaptive gait tasks and (2) while considering the relation of the passive dynamic stability to the overall gait stability. First, authors have interpreted MOS_AP_ depending on the task. For example, Bosse et al. consider XcoM ahead of the BOS boundary as unstable gait while descending stairs [6], whereas others (including us) assume roughly the opposite for obstacle crossing tasks [5, 13]. It seems that these interpretations must be evaluated on a case-by-case basis. A better approach would be to validate the interpretations by correlating MOS_AP_ characteristics with fall rates. Although this is difficult to accomplish, and many biomechanical variables used to quantify stability of human gait lack such validation [54], this is an important goal for future research.

Second, MOS_AP_ captures only the passive dynamic stability of the body, and the relation of the passive stability to the overall stability of human gait is not straightforward due to the contributions of active neuromuscular processes. For example, when MOS_AP_ is positive, indicating that a passive walker would stall due to insufficient energy, higher active push-off by the rear leg of a human walker could overcome this deficit. This is reflected in the data. Although the relationship MOS_AP_ < 0 is consistently observed in stable human gait [7–9, 13, 55–57], the reverse relationship (MOS_AP_ > 0) has also been observed in some stable human walking trials [55]. Even in our data (Fig 3A), we observe positive and negative MOS_AP_ values, and there were no signs of instability in the gait overall.

Nevertheless, the accumulated evidence regarding the changes in MOS_AP_ across tasks and populations suggests that the passive dynamic stability might influence the ability of the person to recover from large perturbations, given that the speed and magnitude of human neuromuscular responses are bounded. Hence passive stability is enhanced when large perturbations could occur. This effect will likely be larger in older or patient populations where maximal capacities are reduced. Therefore, we suggest that studying MOS_AP_ is useful; our study of MOS_AP_ has revealed information regarding locomotor control, and our findings will serve as a baseline for identifying potential locomotor issues in various populations.

Another limitation of this study is that we estimated the CoM location using four pelvis markers [26]. A rigorous whole-body model could have provided slightly different estimates of the CoM motion. However, these differences would influence MOS_AP_ similarly across all conditions, and using a different method to obtain CoM kinematics would likely yield similar qualitative results and overall conclusions, especially given the large effect sizes for all our outcome variables.

It can be argued that the all the steps of the unobstructed task are similar. Therefore, the data for the unobstructed task could be collapsed across the steps and compared with the steps of the obstructed task. We performed this alternate analysis and obtained results that led to identical conclusions. However, we observed effects of gait initiation and termination in our data (cf. Fig 3C). Gait was initiated one step before the examined steps, and gait was terminated one or two steps after the trail crossing step. Therefore, collapsing the data across all steps of the unobstructed task would be suspect, and the statistical approach we used is more appropriate. Using a longer walkway – with more steps before and after the obstacle – may alter the values of our outcome variables. We do not expect that these changes will influence our key conclusions (proactive changes in and control of MOS_AP_). However, effects of gait initiation and termination on MOS_AP_ characteristics may be worth an independent investigation.

## 5. Conclusion

We demonstrated that the XcoM and the step length covary to maintain MOS_AP_, indicating that MOS_AP_ is controlled during unobstructed and obstructed gait. In conjunction with the consistently low MOS_AP_ values for most of the analyzed steps, we conclude that the MOS_AP_ is controlled to exploit the passive dynamics and achieve forward progression at low energetic cost. Furthermore, the value of MOS_AP_ is proactively altered while approaching an obstacle, and MOS_AP_ shows substantial fluctuations for the two crossing steps. These changes reflect a tradeoff between stability and energy efficiency. The changes during approach and lead crossing steps indicate increasingly cautious gait with a growing preference for stability, whereas the opposite change after the trail crossing step indicates a reversion to improving efficiency when the risk to stability is reduced. Thus, our results indicate that humans exploit the passive AP body motion to meet specific ends dictated by the locomotor task. We conclude that the UCM analysis of MOS_AP_ provides new information regarding the control of stability during walking, especially for gait tasks requiring proactive adaptations, and our methods could be valuable in understanding the effects of age and pathology on gait.

## Notes

https://purr.purdue.edu/publications/4150/1

### Competing Interest Statement

The authors have declared no competing interest.

### Summary of Updates

Introduction altered to reflect an updated interpretation of MOSap. Some issues in interpreting MOSap are highlighted in the Limitations section. Order of presentation of results is changed. Alternate statistical approach mentioned in the Limitations.

## REFERENCES

1. King DL, Arnold AS, Smith SL. A Kinematic Comparison of Single, Double and Triple Axels. Journal of Applied Biomechanics. 1994;10(1):51–60. doi: DOI 10.1123/jab.10.1.51. PubMed PMID: WOS:A1994MU82100005.

2. Knoll K, Hildebrand F. Optimum Movement Coordunation in Multi-Revolution Jumps in Figure Skating. In: Riehle HJ, Vieten MM, editors. 16 International Symposium on Biomechanics in Sport July 21–25; Konstanz, Germany 1998.

3. Kuo AD, Donelan JM. Dynamic principles of gait and their clinical implications. Phys Ther. 2010;90(2):157–174. Epub 2009/12/22. doi: 10.2522/ptj.20090125. PubMed PMID: 20023002; PubMed Central PMCID: PMCPMC2816028.

4. Kuo AD, Donelan JM, Ruina A. Energetic consequences of walking like an inverted pendulum: step-to-step transitions. Exerc Sport Sci Rev. 2005;33(2):88–97. Epub 2005/04/12. doi: 10.1097/00003677-200504000-00006. PubMed PMID: 15821430.

5. Hak L, Hettinga FJ, Duffy KR, Jackson J, Sandercock GRH, Taylor MJD. The concept of margins of stability can be used to better understand a change in obstacle crossing strategy with an increase in age. J Biomech. 2019;84:147–152. Epub 2019/01/16. doi: 10.1016/j.jbiomech.2018.12.037. PubMed PMID: 30642664.

6. Bosse I, Oberlander KD, Savelberg HH, Meijer K, Bruggemann GP, Karamanidis K. Dynamic stability control in younger and older adults during stair descent. Hum Mov Sci. 2012;31(6):1560–1570. Epub 2012/08/03. doi: 10.1016/j.humov.2012.05.003. PubMed PMID: 22853941.

7. Bierbaum S, Peper A, Karamanidis K, Arampatzis A. Adaptive feedback potential in dynamic stability during disturbed walking in the elderly. J Biomech. 2011;44(10):1921–1926. Epub 2011/05/11. doi: 10.1016/j.jbiomech.2011.04.027. PubMed PMID: 21555126.

8. Bierbaum S, Peper A, Karamanidis K, Arampatzis A. Adaptational responses in dynamic stability during disturbed walking in the elderly. J Biomech. 2010;43(12):2362–2368. Epub 2010/05/18. doi: 10.1016/j.jbiomech.2010.04.025. PubMed PMID: 20472240.

9. Ohtsu H, Yoshida S, Minamisawa T, Katagiri N, Yamaguchi T, Takahashi T, et al. Does the balance strategy during walking in elderly persons show an association with fall risk assessment? J Biomech. 2020;103:109657. Epub 2020/02/10. doi: 10.1016/j.jbiomech.2020.109657. PubMed PMID: 32035661.

10. Peebles AT, Bruetsch AP, Lynch SG, Huisinga JM. Dynamic Balance Is Related to Physiological Impairments in Persons With Multiple Sclerosis. Arch Phys Med Rehabil. 2018;99(10):2030–2037. Epub 2017/12/25. doi: 10.1016/j.apmr.2017.11.010. PubMed PMID: 29274726; PubMed Central PMCID: PMCPMC6014868.

11. Peebles AT, Bruetsch AP, Lynch SG, Huisinga JM. Dynamic balance in persons with multiple sclerosis who have a falls history is altered compared to non-fallers and to healthy controls. J Biomech. 2017;63:158–163. Epub 2017/09/12. doi: 10.1016/j.jbiomech.2017.08.023. PubMed PMID: 28889946.

12. Peebles AT, Reinholdt A, Bruetsch AP, Lynch SG, Huisinga JM. Dynamic margin of stability during gait is altered in persons with multiple sclerosis. J Biomech. 2016;49(16):3949–3955. Epub 2016/11/28. doi: 10.1016/j.jbiomech.2016.11.009. PubMed PMID: 27889188; PubMed Central PMCID: PMCPMC5176013.

13. Raffegeau TE, Brinkerhoff SA, Kellaher GK, Baudendistel S, Terza MJ, Roper JA, et al. Changes to margins of stability from walking to obstacle crossing in older adults while walking fast and with a dual-task. Exp Gerontol. 2022;161:111710. Epub 2022/01/30. doi: 10.1016/j.exger.2022.111710. PubMed PMID: 35090973.

14. Hof AL, Gazendam MG, Sinke WE. The condition for dynamic stability. J Biomech. 2005;38(1):1–8. Epub 2004/11/03. doi: 10.1016/j.jbiomech.2004.03.025. PubMed PMID: 15519333.

15. Hof AL. The ‘extrapolated center of mass’ concept suggests a simple control of balance in walking. Hum Mov Sci. 2008;27(1):112–125. Epub 2007/10/16. doi: 10.1016/j.humov.2007.08.003. PubMed PMID: 17935808.

16. Tillman M, Ambike S. The Influence of Recent Actions and Anticipated Actions on the Stability of Finger Forces During a Tracking Task. Motor Control. 2020;24(3):365–382. Epub 2020/07/15. doi: 10.1123/mc.2019-0124. PubMed PMID: 32663389.

17. Hof AL, Vermerris SM, Gjaltema WA. Balance responses to lateral perturbations in human treadmill walking. J Exp Biol. 2010;213(Pt 15):2655–2664. Epub 2010/07/20. doi: 10.1242/jeb.042572. PubMed PMID: 20639427.

18. McAndrew Young PM, Wilken JM, Dingwell JB. Dynamic margins of stability during human walking in destabilizing environments. J Biomech. 2012;45(6):1053–1059. Epub 2012/02/14. doi: 10.1016/j.jbiomech.2011.12.027. PubMed PMID: 22326059; PubMed Central PMCID: PMCPMC3321251.

19. Scholz JP, Schoner G. The uncontrolled manifold concept: identifying control variables for a functional task. Exp Brain Res. 1999;126(3):289–306. pubMed PMID: 10382616.

20. Chou LS, Draganich LF, Song SM. Minimum energy trajectories of the swing ankle when stepping over obstacles of different heights. J Biomech. 1997;30(2):115–120. Epub 1997/02/01. doi: 10.1016/s0021-9290(96)00111-x. PubMed PMID: 9001931.

21. Haefeli J, Vogeli S, Michel J, Dietz V. Preparation and performance of obstacle steps: interaction between brain and spinal neuronal activity. Eur J Neurosci. 2011;33(2):338–348. Epub 2010/11/13. doi: 10.1111/j.1460-9568.2010.07494.x. PubMed PMID: 21070395.

22. Harris CM, Wolpert DM. Signal-dependent noise determines motor planning. Nature. 1998;394(6695):780–784. Epub 1998/09/02. doi: 10.1038/29528. PubMed PMID: 9723616.

23. Elias LJ, Bryden MP, Bulman-Fleming MB. Footedness is a better predictor than is handedness of emotional lateralization. Neuropsychologia. 1998;36(1):37–43. Epub 1998/04/09. doi: 10.1016/s0028-3932(97)00107-3. PubMed PMID: 9533385.

24. Rosenblatt NJ, Hurt CP. Recommendation for the minimum number of steps to analyze when performing the uncontrolled manifold analysis on walking data. J Biomech. 2019;85:218–223. Epub 2019/02/06. doi: 10.1016/j.jbiomech.2019.01.018. PubMed PMID: 30718066; PubMed Central PMCID: PMCPMC6905614.

25. Desailly E, Daniel Y, Sardain P, Lacouture P. Foot contact event detection using kinematic data in cerebral palsy children and normal adults gait. Gait Posture. 2009;29(1):76–80. Epub 2008/08/05. doi: 10.1016/j.gaitpost.2008.06.009. PubMed PMID: 18676147.

26. Whittle MW. Three-dimensional motion of the center of gravity of the body during walking. Hum Movement Sci. 1997;16(2-3):347–355. doi: Doi 10.1016/S0167-9457(96)00052-8. PubMed PMID: WOS:A1997WV17500014.

27. Hak L, Houdijk H, Beek PJ, van Dieen JH. Steps to take to enhance gait stability: the effect of stride frequency, stride length, and walking speed on local dynamic stability and margins of stability. PLoS One. 2013;8(12):e82842. Epub 2013/12/19. doi: 10.1371/journal.pone.0082842. PubMed PMID: 24349379; PubMed Central PMCID: PMCPMC3862734.

28. Cui C, Kulkarni A, Rietdyk S, Barbieri FA, Ambike S. Synergies in the ground reaction forces and moments during double support in curb negotiation in young and older adults. J Biomech. 2020;106:109837. Epub 2020/06/11. doi: 10.1016/j.jbiomech.2020.109837. PubMed PMID: 32517974.

29. Kulkarni A, Cho H, Rietdyk S, Ambike S. Step length synergy is weaker in older adults during obstacle crossing. J Biomech. 2021;118:110311. Epub 2021/02/19. doi: 10.1016/j.jbiomech.2021.110311. PubMed PMID: 33601182.

30. Vaz DV, Pinto VA, Junior RRS, Mattos DJS, Mitra S. Coordination in adults with neurological impairment - A systematic review of uncontrolled manifold studies. Gait Posture. 2019;69:66–78. Epub 2019/01/25. doi: 10.1016/j.gaitpost.2019.01.003. PubMed PMID: 30677709.

31. Latash ML, Huang X. Neural control of movement stability: Lessons from studies of neurological patients. Neuroscience. 2015;301:39–48. doi: 10.1016/j.neuroscience.2015.05.075. PubMed PMID: 26047732; PubMed Central PMCID: PMCPMC4504804.

32. Ambike S, Penedo T, Kulkarni A, Santinelli FB, Barbieri FA. Step length synergy while crossing obstacles is weaker in patients with Parkinson’s disease. Gait Posture. 2021;84:340–345. Epub 2021/01/18. doi: 10.1016/j.gaitpost.2021.01.002. PubMed PMID: 33454501.

33. Sivakumaran S, Schinkel-Ivy A, Masani K, Mansfield A. Relationship between margin of stability and deviations in spatiotemporal gait features in healthy young adults. Hum Mov Sci. 2018;57:366–373. Epub 2017/10/11. doi: 10.1016/j.humov.2017.09.014. PubMed PMID: 28987772; PubMed Central PMCID: PMCPMC5770210.

34. Worden TA, Vallis LA. Stability control during the performance of a simultaneous obstacle avoidance and auditory Stroop task. Exp Brain Res. 2016;234(2):387–396. Epub 2015/10/22. doi: 10.1007/s00221-015-4461-z. PubMed PMID: 26487180.

35. Jin J, van Dieen JH, Kistemaker D, Daffertshofer A, Bruijn SM. Does ankle push-off correct for errors in anterior-posterior foot placement relative to center-of-mass states? BioRxiv. 2022. doi: https://doi.org/10.1101/2022.03.14.484283.

36. Kuo AD. A simple model of bipedal walking predicts the preferred speed-step length relationship. J Biomech Eng. 2001;123(3):264–269. Epub 2001/07/31. doi: 10.1115/1.1372322. PubMed PMID: 11476370.

37. Chou LS, Draganich LF. Stepping over an obstacle increases the motions and moments of the joints of the trailing limb in young adults. J Biomech. 1997;30(4):331–337. Epub 1997/04/01. doi: 10.1016/s0021-9290(96)00161-3. PubMed PMID: 9075000.

38. Winter DA. The Bionechanics and Motor Control of Human Gait: Normal, Elderly and Pathological. Second ed. Kitchener, ON: Waterloo Biomechanics; 1991.

39. Dietz V, Quintern J, Sillem M. Stumbling reactions in man: significance of proprioceptive and pre-programmed mechanisms. J Physiol. 1987;386:149–163. Epub 1987/05/01. doi: 10.1113/jphysiol.1987.sp016527. PubMed PMID: 3681704; PubMed Central PMCID: PMCPMC1192455.

40. Takakusaki K. Neurophysiology of gait: from the spinal cord to the frontal lobe. Mov Disord. 2013;28(11):1483–1491. Epub 2013/10/18. doi: 10.1002/mds.25669. PubMed PMID: 24132836.

41. Feldman AG, Levin MF, Garofolini A, Piscitelli D, Zhang L. Central pattern generator and human locomotion in the context of referent control of motor actions. Clin Neurophysiol. 2021;132(11):2870–2889. Epub 2021/10/11. doi: 10.1016/j.clinph.2021.08.016. PubMed PMID: 34628342.

42. Latash ML. One more time about motor (and non-motor) synergies. Exp Brain Res. 2021;239(10):2951–2967. Epub 2021/08/13. doi: 10.1007/s00221-021-06188-4. PubMed PMID: 34383080.

43. Bauby CE, Kuo AD. Active control of lateral balance in human walking. J Biomech. 2000;33(11):1433–1440. Epub 2000/08/15. doi: 10.1016/s0021-9290(00)00101-9. PubMed PMID: 10940402.

44. Collins SH, Kuo AD. Two independent contributions to step variability during over-ground human walking. PLoS One. 2013;8(8):e73597. Epub 2013/09/10. doi: 10.1371/journal.pone.0073597. PubMed PMID: 24015308; PubMed Central PMCID: PMCPMC3756042.

45. Patla AE, Vickers JN. Where and when do we look as we approach and step over an obstacle in the travel path? Neuroreport. 1997;8(17):3661–3665. Epub 1998/01/14. doi: 10.1097/00001756-199712010-00002. PubMed PMID: 9427347.

46. Diaz GJ, Parade MS, Barton SL, Fajen BR. The pickup of visual information about size and location during approach to an obstacle. PLoS One. 2018;13(2):e0192044. Epub 2018/02/06. doi: 10.1371/journal.pone.0192044. PubMed PMID: 29401511; PubMed Central PMCID: PMCPMC5798835.

47. Tillman M, Ambike S. Cue-induced changes in the stability of finger force-production tasks revealed by the uncontrolled manifold analysis. J Neurophysiol. 2018;119(1):21–32. Epub 2017/09/22. doi: 10.1152/jn.00519.2017. PubMed PMID: 28931612.

48. McAndrew Young PM, Dingwell JB. Voluntarily changing step length or step width affects dynamic stability of human walking. Gait Posture. 2012;35(3):472–477. Epub 2011/12/17. doi: 10.1016/j.gaitpost.2011.11.010. PubMed PMID: 22172233; PubMed Central PMCID: PMCPMC3299923.

49. van Leeuwen AM, van Dieen JH, Daffertshofer A, Bruijn SM. Active foot placement control ensures stable gait: Effect of constraints on foot placement and ankle moments. PLoS One. 2020;15(12):e0242215. Epub 2020/12/18. doi: 10.1371/journal.pone.0242215. PubMed PMID: 33332421; PubMed Central PMCID: PMCPMC7746185.

50. van Leeuwen AM, van Dieen JH, Daffertshofer A, Bruijn SM. Ankle muscles drive mediolateral center of pressure control to ensure stable steady state gait. Sci Rep. 2021;11(1):21481. Epub 2021/11/04. doi: 10.1038/s41598-021-00463-8. PubMed PMID: 34728667; PubMed Central PMCID: PMCPMC8563802.

51. Rosenblatt NJ, Grabiner MD. Measures of frontal plane stability during treadmill and overground walking. Gait Posture. 2010;31(3):380–384. Epub 2010/02/05. doi: 10.1016/j.gaitpost.2010.01.002. PubMed PMID: 20129786.

52. Reimann H, Fettrow T, Jeka JJ. Strategies for the Control of Balance During Locomotion. Kinesiology Review. 2018;7(1):18. doi: 10.1123/kr.2017-005310.1123/kr.2017-005310.1123/kr.2017-005310.1123/kr.2017-0053.

53. Kazanski ME, Cusumano JP, Dingwell JB. Rethinking margin of stability: Incorporating step-to-step regulation to resolve the paradox. J Biomech. 2022;144:111334. Epub 2022/10/17. doi: 10.1016/j.jbiomech.2022.111334. PubMed PMID: 36244320.

54. Bruijn SM, Meijer OG, Beek PJ, van Dieen JH. Assessing the stability of human locomotion: a review of current measures. J R Soc Interface. 2013;10(83):20120999. Epub 2013/03/22. doi: 10.1098/rsif.2012.0999. PubMed PMID: 23516062; PubMed Central PMCID: PMCPMC3645408.

55. Hohne A, Stark C, Bruggemann GP, Arampatzis A. Effects of reduced plantar cutaneous afferent feedback on locomotor adjustments in dynamic stability during perturbed walking. J Biomech. 2011;44(12):2194–2200. Epub 2011/07/06. doi: 10.1016/j.jbiomech.2011.06.012. PubMed PMID: 21726865.

56. Ohtsu H, Yoshida S, Minamisawa T, Takahashi T, Yomogida SI, Kanzaki H. Investigation of balance strategy over gait cycle based on margin of stability. J Biomech. 2019;95:109319. Epub 2019/08/31. doi: 10.1016/j.jbiomech.2019.109319. PubMed PMID: 31466715.

57. Watson F, Fino PC, Thornton M, Heracleous C, Loureiro R, Leong JJH. Use of the margin of stability to quantify stability in pathologic gait - a qualitative systematic review. Bmc Musculoskel Dis. 2021;22(1). doi: 10.1186/s12891-021-04466-4. PubMed PMID: WOS:000669563500004.

